# Periventricular nodular heterotopia is functionally coupled with the neocortex during resting and task states

**DOI:** 10.1101/2022.03.03.482573

**Authors:** Yayue Gao, Guanpeng Chen, Pengfei Teng, Xin Zhang, Fang Fang, Dario J. Englot, Guoming Luan, Xiongfei Wang, Qian Wang

## Abstract

Periventricular nodular heterotopia (PVNH) is a well-defined developmental disorder characterized by failed neuronal migration, which forms ectopic neuronal nodules along the ventricular walls. Previous studies mainly focus on clinical symptoms caused by the PVNH tissue, such as seizures. However, little is known about whether and how neurons in the PVNH tissue functionally communicate with neurons in the neocortex. To probe this, we applied magnetoencephalography (MEG) and stereo-electroencephalography (sEEG) recordings to patients with PVNH during resting and task states. By estimating frequency-resolved phase coupling strength of the source-reconstructed neural activities, we found that the PVNH tissue was spontaneously coupled with the neocortex in the α to β frequency range, which was consistent with the synchronization pattern within the neocortical network. Furthermore, the coupling strength between PVNH and sensory areas effectively modulated the local neural activity in sensory areas. In both MEG and sEEG visual experiments, the PVNH tissue exhibited visual evoked responses, with a similar pattern and latency as the ipsilateral visual cortex. These findings demonstrate that PVNH is functionally integrated into cognition-related cortical circuits, suggesting a co-development perspective of ectopic neurons after their migration failure.

## Introduction

In the developing brain, neurons migrate from areas they are born to areas they will settle, constructing the neocortex.^1–3^ However, abnormal neuronal migration leads to malformations, such as periventricular nodular heterotopia (PVNH), which is characterized by the stranded nodular gray matter around the lateral ventricular walls.^2,4–6^ Although the PVNH tissue may become involved in generating seizures,^7–11^ recent functional magnetic resonance imaging (fMRI) studies suggested a functional role of the PVNH tissue by finding interactions between the PVNH tissue and the neocortex.^9,12–16^

However, electrophysiological evidence supporting the functional role of the PVNH tissue is rare. In a recent study, using stereo-electroencephalography (sEEG) recordings, it was shown that PVNH tissues not only exhibited spontaneous activities which were correlated to the neocortex, but also generated normal physiological responses during a cognitive task in three patients.^17^ Although this landmark study suggested a functional involvement of the PVNH tissue, due to the sparseness of sEEG sampling, we are still unclear whether and how neurons in the PVNH tissue functionally communicate with neurons in the neocortex over a wide spatial range, which limits our understanding of the potential cognitive relevance of PVNH neurons.

To fill this gap, here we report a quantitative magnetoencephalography (MEG) study of patients with PVNH. We reconstructed spontaneous neural activities in the PVNH tissue and brain-wide areas with high spatial, temporal, and frequency resolution and estimated the inter-areal coupling strength using phase locking values (PLVs).^18,19^ We then inspected whether local activities in sensory areas were modulated by their communications with PVNH tissues. In addition, in rare opportunities, we compared visual evoked responses in the PVNH tissue and the visual cortex in two patients, using MEG reconstructed signals and sEEG signals, respectively.

## Methods

### Patients

From May 2014 to November 2020, thirteen patients with PVNH were recruited from the clinical database of epilepsy patients at Sanbo Brain Hospital, Capital Medical University, including six patients with left PVNH tissues, six patients with right PVNH tissues, and one patient with bilateral PVNH tissues (Figure 1A). The detailed demographic and clinical information of each patient was shown in Table 1.

**Table 1.** Demographic and clinical information of patients with periventricular nodular heterotopia

| Patient No. | Gender | Age | PVNH | Age at seizure onset (years) | Epilepsy duration (years) | Seizure frequency (per month) |
| --- | --- | --- | --- | --- | --- | --- |
| 1 | M | 26 | L | 23 | 3 | 10 |
| 2 | M | 22 | L | 20 | 2 | 0.25 |
| 3 | F | 48 | L | 18 | 30 | 3 |
| 4 | F | 38 | L | 12 | 26 | 1 |
| 5 | F | 51 | L | 30 | 21 | 1 |
| 6 | F | 30 | L | 27 | 3 | 0.5 |
| 7 | M | 19 | R | 12 | 7 | 0.25 |
| 8 | M | 13 | R | 7 | 6 | 3 |
| 9 | F | 13 | R | 1 | 12 | 2 |
| 10 | F | 22 | R | 19 | 3 | 1 |
| 11 | M | 18 | R | 15 | 3 | 3 |
| 12 | M | 37 | R | 36 | 0.9 | 0.5 |
| 13 | F | 23 | B | 23 | 0.2 | 3 |

F, female; M, male; L, left; R, right; B, bilateral.

**Figure 1.**
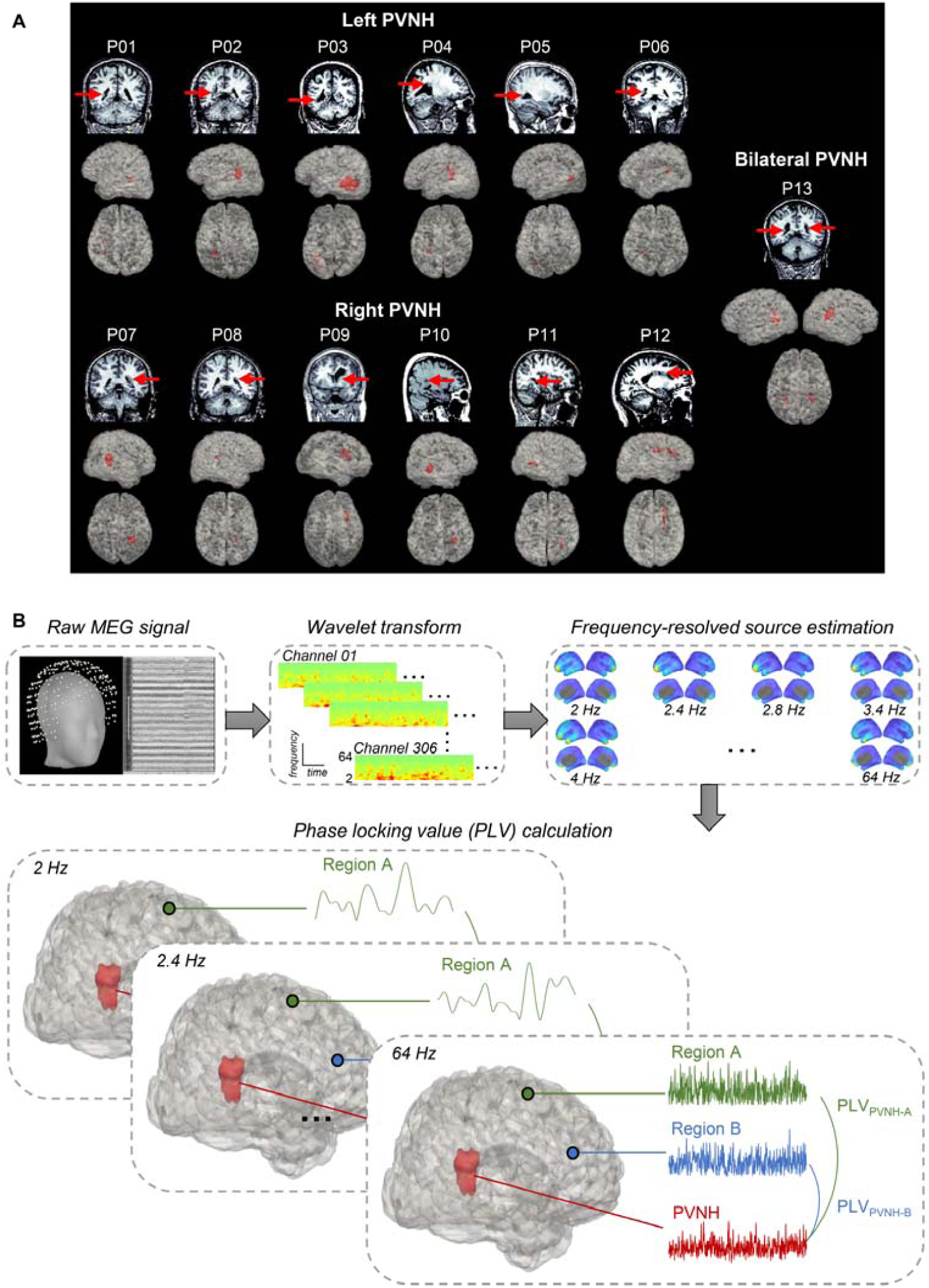
PVNH localizations and MEG data analysis procedure. (**A**) Localizations of PVNH in 13 patients are exhibited in anatomical MR images (T1-weighted) and 3D reconstructed brain models (in lateral and top views), respectively. Red arrows and areas indicate PVNH tissues. (**B**) A schematic illustration of the MEG data analysis procedure. The continuous MEG signals were firstly decomposed to spectra-temporal components using Morlet wavelet transforms, logarithmically ranging from 2 to 64 Hz. Then, the source estimation was conducted within each frequency. Inter-areal phase-locking values (PLVs) were calculated between the PVNH tissue and each cortical area in each frequency.

### Standard Protocol Approvals, Registrations, and Patient Consents

All patients provided written informed consent in accordance with research protocols approved by the Ethics Committee of the Sanbo Brain Hospital, Capital Medical University.

### MEG Recording and Pre-Processing

Neuromagnetic signals were recorded using a 306-sensor (204 planar gradiometers, 102 magnetometers) whole-head MEG system (Elekta-Neuromag, Helsinki, Finland), sampled at 1000 Hz. The head position inside MEG was determined by four head position indicator (HPI) coils. The positions of three anatomical landmarks (nasion, left, and right pre-auricular points), four HPI coils, and at least 150 points on the scalp were digitized before MEG recordings. For each patient, five sequences of six-minute resting-state MEG signals were recorded while the patient was lying in a magnetically shielded recording room. In one patient (P11), two blocks of task-state signals were also recorded.

For MEG source localization purposes, a structural T1-weighted MRI dataset (voxel size: 1×0.5×0.5 mm^3^) was acquired for each patient using a 1.5 T Philips Achieva (Best, The Netherlands) MRI scanner. The cortical reconstruction and segmentation were computed to extract the brain volume, cortex surface, and innermost skull surface based on individual T1-weighted images using the Brainsuite software (http://brainsuite.org/). The whole brain was subsequently parcellated into 130 neocortical areas (65 areas in each hemisphere) according to the USCBrain atlas.^20^

The external interference in raw signals was removed by the temporal signal space separation (tSSS) method and then the signals were down-sampled to 400 Hz. Fifty-hertz line-noise and its harmonics were excluded with a band-stop finite impose response (FIR) filter with a band-stop width of 1 Hz. For each MEG sensor, the continuous signal was then decomposed to spectra-temporal components (Figure 1B) using Morlet wavelet transforms by *morlet_transform* function in Brainstorm.^21^ The center frequencies of spectra-temporal components were organized in log space, from 2 to 64 Hz.

### Frequency-Resolved Source Estimation

For the continuous signal in each frequency, we performed two types of source estimation. Surface-based and volume-based models were applied to obtain the reconstructed MEG time series in 130 neocortical areas and the PVNH tissue, respectively. All procedures were performed on the individual brain model using the Brainstorm toolbox.

For the surface-based source estimation, we computed individual MEG forward models using the overlapping-sphere method on the individual pial surface (~15,000 vertices). Then, using the weighted minimum norm estimate (wMNE) model, we obtained source-reconstructed MEG time series on 130 neocortical areas.

For the volume-based source estimation, we also conducted the overlapping-sphere method on the individual volume space. Similarly, we obtained volume-based MEG time series by the wMNE method. Voxels within the PVNH tissue were identified manually by two experienced neurosurgeons (Dr. Xiongfei Wang and Dr. Pengfei Teng). In each PVNH tissue, we averaged the time series in all PVNH voxels for further analyses.

### Inter-Areal PLVs

For source-reconstructed time series in the neocortex and PVNH, we estimated inter-areal phase interactions at the individual level using PLVs. For each frequency, the complex PLV (cPLV) between two time series, *x*(*t*) and *y*(*t*), is computed as:

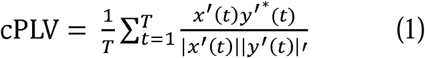

where *T* is the total number of time points of the entire signal, and * is complex conjugate, *x*′(*t*) and *y*′ (*t*) are the complex wavelet coefficients of *x*(*t*) and *y*(*t*), respectively. To reduce the influence of the volume conduction effect, the PLV was defined as the absolute imaginary part of cPLV (PLV = |Im(cPLV)|), a metric insensitive to zero-lag phase interactions.^18,19^ The PLV is a scalar measure bounded between 0 and 1.

We then grouped and averaged PLVs in five frequency bands (δ, 2-4 Hz; θ, 4 - 8 Hz; α, 8 - 16 Hz; β, 16 - 32 Hz; and γ, 32 - 64 Hz), respectively. We defined the PVNH-cortical PLVs as the PLVs between PVNH tissues and neocortical areas and the cortico-cortical PLVs as the PLVs among neocortical areas. And the relative PLV equals the absolute PLV minus the surrogate mean (**see Statistical Analyses**).

### Comparison of PSDs Between Different PLVs

To test whether the changes of local neural activities were correlated with PLVs between the PVNH tissue and a certain neocortical area, we compared the local power spectral densities (PSDs) of source-reconstructed time series with high PVNH-cortical PLVs and those with low PVNH-cortical PLVs in each frequency band. The averaged source-reconstructed signals in auditory, visual, and somatosensory areas were first partitioned into several 20-second segments. For each neocortical area, ten segments with the highest PLVs (high-PLV group) and ten with the lowest PLVs (low-PLV group) were selected in each frequency band. After Fourier transform, we computed and averaged PSDs of time segments in the high-PLV group (*PSD*_*high*_) and those in the low-PLV group (*PSD*_*low*_). The normalized PSD difference was calculated as follows:

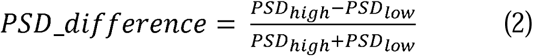

### Procedure and Data Analyses in the MEG Visual Experiment

One patient (P11) participated in the MEG visual experiment. Visual stimuli were projected onto a translucent screen via a DLP projector (refresh rate: 60 Hz; spatial resolution: 1280×1024; PT-D7700E-K). The patient was seated in a magnetically shielded recording room and the viewing distance was 85 cm.

A flickering full-contrast checkerboard stimulus (size: 17°× 25° ; Michelson contrast: 1; mean luminance: 288.83 cd/m^2^) was presented in the left or right visual hemifield against a gray background (luminance: 35.4 cd/m^2^). The checkerboard stimulus reversed in phase every 500 ms, with a total of 200 reversals in each block. During the experiment, two blocks, one for each hemifield, were completed. The patient was asked to fixate at the fixation point.

The raw electrophysiological signals were first imported into the Brainstorm toolbox. Both volume-based and surface-based source estimations were applied using the same parameters in the resting-state experiment. We conducted the volume-based source estimation, identified the voxels within the PVNH tissue, and averaged the time series in all PVNH voxels for further analyses. We also applied the surface-based source estimation to reconstruct neural activities in visual cortex. We defined visual cortex as 100 vertices with the largest M100 (~95 ms) visual evoked fields in the hemisphere ipsilateral to the PVNH tissue. Continuous source-reconstructed signals in the PVNH tissue and visual cortex were epoched starting at 100 ms before stimulus reversal and ending at 400 ms after stimulus reversal. Visual evoked fields (absolute value) were computed by averaging the signals across trials. For each epoch, the latency was defined as the time point of the first peak after 50 ms.

### Procedure and Data Analyses in the SEEG Visual Experiment

One patient (P04) performed the visual task during stereo-electroencephalography (sEEG) recordings. The patient was implanted with several stereo-electrodes, which contained 8-16 independent contacts (0.8 mm in diameter, 2 mm in length, intercontact spacing of 3.5 mm). To identify the location of each contact, we co-registered the post-implantation CT with the pre-implantation T1-weighted MR images and manually labelled each electrode on the aligned CT images. The contacts within the PVNH tissue and visual cortex were confirmed by two experienced neurosurgeons (Dr. Xiongfei Wang and Dr. Pengfei Teng).

The sEEG recordings were conducted in the ward after electrode implantation, with a sampling rate at 512 Hz (Nicolet Clinical Amplifier, USA). Visual stimuli were displayed on a laptop (refresh rate: 60 Hz; spatial resolution: 1980×1080; 14-inch, Thinkpad T590). The viewing distance was 42 cm, and the patient’s head was stabilized with a chin rest. A full-screen, square-wave grating (radius: 17.8 degrees; spatial frequency: 1 cycle/degree; Michelson contrast: 1; mean luminance: 160.98 cd/m^2^) was presented against a gray background (luminance: 22.4 cd/m^2^). The orientation of the grating could be one of four possible orientations from 0° to 135° in steps of 45° (relative to the vertical axis). In each trial, the stimulus was presented for 500 ms, followed by a 500-ms blank interval. During the experiment, a total of eighty trials, twenty trials for each orientation, were completed in a random order. The patient was asked to fixate at the fixation point and detect its color change by mouse clicking.

The raw electrophysiological signals were first imported into the EEGLAB toolbox^22^ and visually inspected for artifact rejections. Then the signals were bandpass filtered at 0.5 – 200 Hz, epoched starting at 100 ms before stimulus onset and ending at 400 ms after stimulus onset, and corrected for baseline over the 100-ms pre-stimulus baseline. Visual evoked potentials were computed by averaging the signals across trials of all four orientations. For each epoch, the latency was defined as the time point of the first peak after 50 ms.

### Statistical Analyses

To test the statistical significance of PLVs, we estimated the null distributions of interaction metrics with surrogates that preserved the temporal autocorrelation structure of the original signals while eliminating correlations between two signals. In each iteration, for a signal in the PVNH tissue, we divided the whole time series T into two blocks at a random time point *k* so that *x*′_1_(*t*) = *x*′ (1 … *k*) and *x*′_2_(*t*) = *x*′ (*k* … *T*), constructed the surrogate as *x*′_*surr*_ (*t*) = [*x*′_2_, *x*′_1_], and computed the surrogate PLVs as in *Equation (1)*. This randomization procedure was repeated 2,000 times. We then averaged surrogate PLVs and obtained a surrogate mean in each patient for further statistical testing.

Permutation tests were applied to estimate the significance of PLV differences by *bst_permtest* in Brainstorm. Pearson correlation was calculated by *corr* function in MATLAB. Bootstrap procedures (5000 times with replacement) were applied to test the PSD difference against zero and the statistical difference between response latencies. False discovery rate (FDR) corrections were applied to p-values involving multiple comparisons, using the *mafdr* function in MATLAB.

### Data Availability

The data that support the findings of this study are available from the corresponding author, upon reasonable request.

## Results

### PVNH Was Coupled With the Neocortex During the Resting State

At the individual level, we first estimated the spectral property of inter-areal phase synchronization between the PVNH tissue and neocortical areas using source-reconstructed resting-state MEG signals. Generally, as shown in Figure 2A, both the averaged PVNH-cortical and cortico-cortical PLVs were greater than the surrogate means in a broad frequency scale (from 2.4 to 38 Hz, permutation tests, all *p* < 0.005, FDR corrected). As shown in whole-brain statistical maps in Figure 2C and D, after combining PLVs into five frequency bands (δ, 2 - 4 Hz; θ, 4 - 8 Hz; α, 8 - 16 Hz; β, 16 - 32 Hz; and γ, 32 - 64 Hz), we identified that both PVNH-cortical and cortico-cortical PLV maps exhibited a peak in the α-β band (permutation tests, all *p* < 0.005, FDR corrected), which was similar to the spectral property of cortico-cortical PLVs in healthy participants.^18^

**Figure 2.**
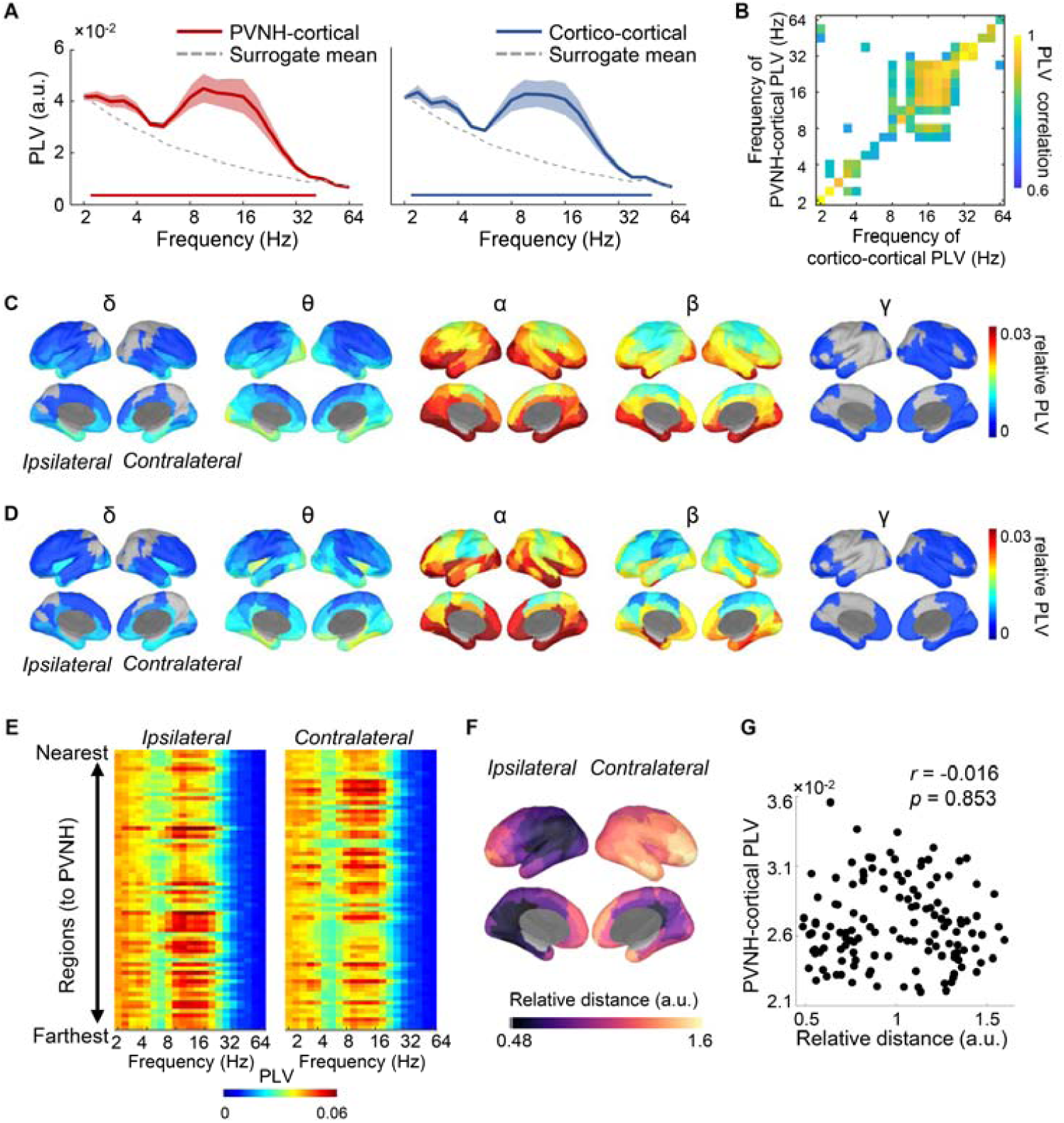
PVNH-cortical and cortico-cortical PLVs. (**A**) Frequency-resolved PVNH-cortical PLVs *(left*) and cortico-cortical PLVs *(right*) are exhibited in coloured lines. Shaded areas exhibit SEM of PLVs. Dash lines are the 95th%-ile of the surrogate mean (determined by bootstrap with 5000 times). Coloured bars near the horizontal axis indicate significance (permutation tests, *p* < 0.005, FDR corrected). (**B**) The correlation map between PVNH-cortical PLVs and cortico-cortical PLVs. Only significant correlations were coloured (Pearson correlation, *p* < 0.005, FDR corrected). (**C**) Relative PVNH-cortical and (**D**) cortico-cortical PLV maps in δ (2 - 4 Hz), θ (4 - 8 Hz), α (8 - 16 Hz), β (16 - 32 Hz), and γ (32 - 64 Hz) bands, respectively. The relative PLV equals the absolute PLV minus the surrogate mean in each frequency band. In each cortical area (parcellated using the USCBrain atlas), the averaged PLVs are re-projected to the hemisphere either ipsilateral or contralateral to the individual PVNH location, respectively. Only areas with a PLV greater than the surrogate mean were coloured (permutation tests, *p* < 0.005, FDR corrected). (**E**) PVNH-cortical PLVs sorted by the PVNH-cortical distance. Each line represents PLVs in one cortical area. (**F**) PVNH-cortical distance maps of the hemispheres ipsilateral and contralateral to the individual PVNH location, respectively. (**G**) No correlation between the PVNH-cortical PLV and the PVNH-cortical distance.

Next, we asked whether the PVNH-cortical and cortico-cortical PLVs shared a similar spectral property. As shown in Figure 2A, we found no significant difference between averaged PVNH-cortical and cortico-cortical PLVs in any frequency band (permutation tests, all *p* > 0.05,). Furthermore, the correlation coefficients between PVNH-cortical and cortico-cortical PLVs were clustered in the α-β band (Figure 2B; Pearson correlation, all *p* < 0.05, FDR corrected). The similarity between PVNH-cortical and cortico-cortical PLVs suggested that PVNH might be involved in the cortico-cortical network.

Considering the whole brain as a volume conductor, the high PVNH-cortical PLV could be an effect of a short distance between the PVNH tissue and a certain cortical area. To test this possibility, we sorted the frequency-resolved PLVs of cortical areas by their PVNH-cortical distances (Figure 2E and F). As shown in Figure 2E, areas near the PVNH tissue did not have stronger PLVs than the other areas in any frequency band. No significant correlation was found between the averaged PVNH-cortical PLV and the PVNH-cortical distance (Figure 2G; Pearson correlation, *r* = −0.016, *p* = 0.853). These results demonstrated that the PVNH-cortical PLV was not linearly correlated to distance.

### PVNH-cortical PLVs Were Associated With Activity Changes in Sensory Areas

In healthy participants, a major feature of the spontaneous phase coupling in sensory areas was a peak in the α-β band.^23^ As shown in Figure 2C, the PVNH tissue was coupled with sensory areas including auditory, visual, and somatosensory areas in the α-β band, suggesting that the PVNH tissue might be integrated into sensory systems. To further test that, we estimated whether local PSDs in sensory areas were associated with their interactions with PVNH tissues. As shown in Figure 3A, source-reconstructed signals in auditory, visual, and somatosensory areas were segmented and sorted by their PVNH-cortical PLVs, respectively. We then calculated the PSDs of time series with the highest PLVs and those with the lowest PLVs and compared the PSD difference with zero. In that case, a significant difference between the PSD difference and zero indicated a co-modulated relationship between the local PSD and the PVNH-cortical PLV in a certain area.

**Figure 3.**
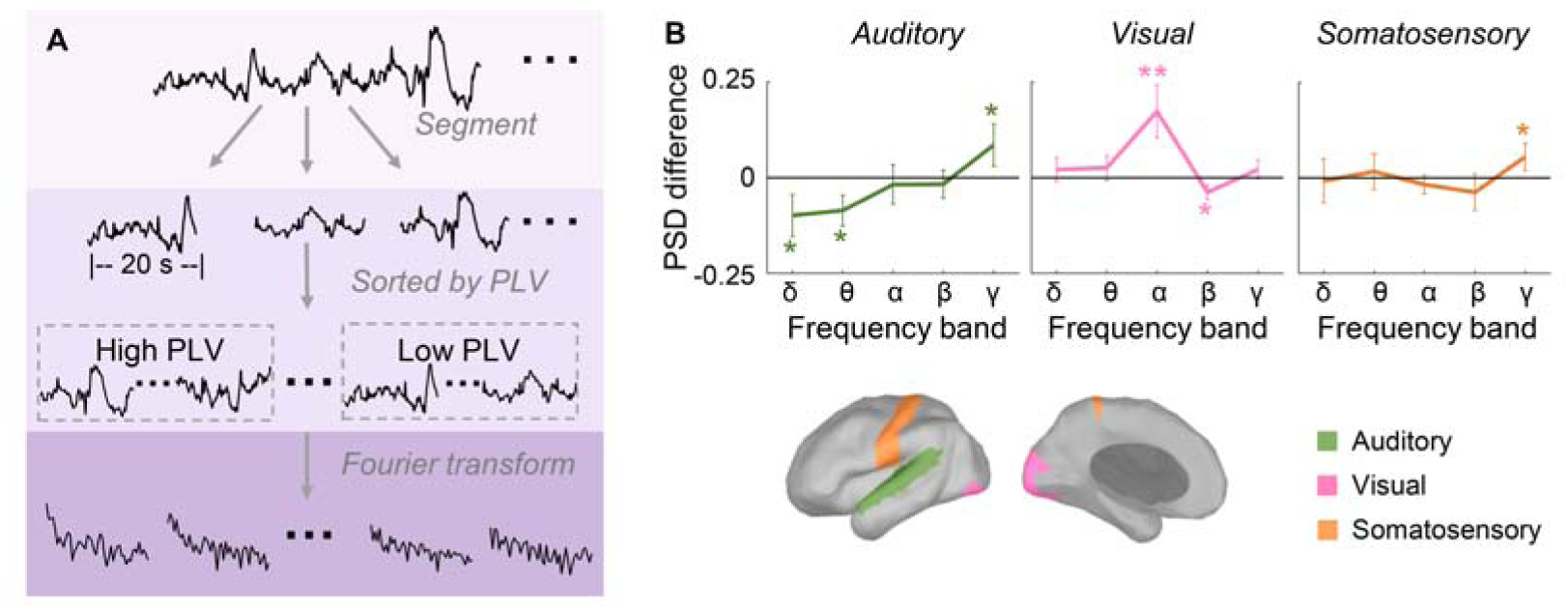
Local PSDs were associated with the PVNH-cortical PLV fluctuations in sensory areas. (**A**) A schematic illustration of PLV-related PSD analysis procedure. Step 1: source-reconstructed signals were partitioned into 20-second segments; Step 2: in each frequency band, segments were sorted by their PLVs, then segments with the highest 10 PLVs (high-PLV group) and the lowest 10 PLVs (low-PLV group) were included in further analyses; Step 3: we calculated the PSDs of those two groups of segments using Fourier transform. (**B**) Frequency-resolved PSD differences between the high-PLV group and the low-PLV group in auditory, visual and somatosensory areas. Error bars indicate SEM. **, *p* < 0.005; *, *p* < 0.05.

As shown in Figure 3B, in auditory cortex, the PSD differences in δ, θ, and γ bands were different from zero (bootstrap tests, δ, 95% confidence interval [CI] −0.15–−0.02, *p* = 0.018; θ, 95% CI −0.15–−0.02, *p* = 0.012; γ, 95% CI 0.01–0.18, *p* = 0.003), which indicated higher PVNH-cortical PLVs were associated with lower PSDs in δ and θ bands and higher PSD in the γ band. In visual cortex, the PSD differences in α and β bands were different from zero (bootstrap tests, α, 95% CI 0.07–0.28, *p* = 0.003; β, 95% CI −0.07–−0.01, *p* = 0.013), demonstrating that the PSD in the α band was enhanced with higher PVNH-cortical PLVs, while the PSD in the β band changed oppositely. In somatosensory cortex, the PSD difference in the γ band was larger than zero (bootstrap test, 95% CI 0.003–0.11, *p* = 0.035), illustrating that larger PLVs led to greater power in the γ band. These results suggested that local PSDs in sensory areas were associated with their interactions with PVNH tissues, with distinct spectral characteristics.

### Visual Evoked Responses in the PVNH Mimicked the Ipsilateral Visual Cortex

Although our results showed that the PVNH tissue was coupled with sensory areas during the resting state, it remained unknown how the PVNH tissue responded to sensory stimuli and communicated with sensory areas during the task state. To explore that, one patient (P11) also participated in a MEG visual experiment. As shown in Figure 4A, visual stimuli were presented to P11 during MEG recordings. We reconstructed the visual evoked activities in both PVNH and the ipsilateral visual cortex. The results showed that both PVNH and visual cortex strongly responded to the visual stimulus presented in the contralateral visual field (Figure 4B). The remarkable visual evoked activity in the PVNH tissue might result from projections from either visual cortex or visual thalamus. To verify the hierarchical relationship between the PVNH tissue and visual cortex, we further compared the first peak latencies of the PVNH tissue and visual cortex at single-trial level, but no significant difference was found (Figure 4E, bootstrap test, *p* > 0.05). These results implied a parallel processing of visual information in PVNH and visual cortex neurons.

**Figure 4.**
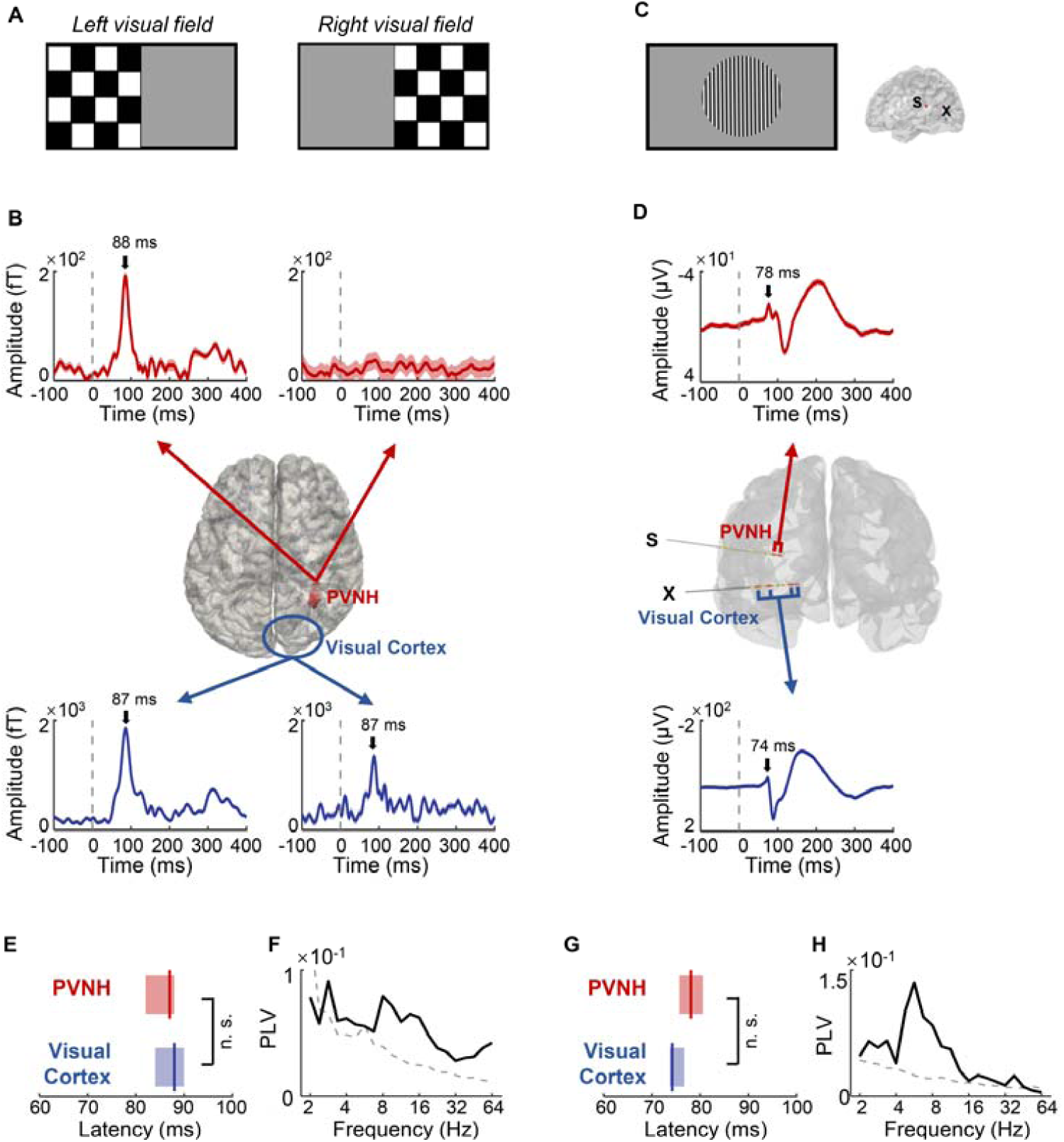
Visual evoked activities in PVNH and visual cortex measured by MEG and sEEG. (**A**) Visual stimuli used in the MEG visual experiment (P11). (**B**) Reconstructed visual evoked fields estimated by source estimation in the PVNH tissue (*red*) and visual cortex (*blue*), respectively. Vertical dashed lines denote the stimulus onset. Black arrows indicate peak latencies. Shaded areas indicate SEM across trials. (**C**) Visual stimuli used in the sEEG visual experiment (P04) *(left*) and stereo-electrodes located in PVNH (Electrode S) and visual cortex (Electrode X) *(right*). (**D**) Visual evoked potentials in the contacts within the PVNH tissue (*red*) and visual cortex (*blue*), respectively. Shaded areas indicate SEM across trials. Comparison between latencies of the PVNH tissue (*red*) and those of visual cortex (*blue*) in the MEG visual experiment (**E**) and the sEEG visual experiment (**G**) are shown. Shaded areas indicate the 5th%-ile to 95th%-ile confidence interval determined by bootstrap with 5000 times. Task-state PLVs between the PVNH tissue and visual cortex in the MEG visual experiment (**F**) and the sEEG visual experiment (**H**) are exhibited. Dash lines are the 95th%-ile of the surrogate mean (determined by bootstrap with 5000 times). *n*.*s*.: not significant.

We further verified this observation via sEEG recordings in a rare opportunity. Another patient (P04) was implanted with stereo-electrodes in both the PVNH tissue and visual cortex for the preoperative evaluation and participated in the sEEG visual experiment. As shown in Figure 4C and D, both contacts located in the PVNH tissue and visual cortex exhibited strong visual evoked potentials. Similar to the MEG visual experiment, we found no significant difference between the latency of PVNH and that of visual cortex (Figure 4G, bootstrap test, *p* > 0.05). These results further confirmed that the functional activity in PVNH might not result from projections from visual cortex, but rather directly from subcortical structures.

We also measured the PLVs between the PVNH tissue and visual cortex during the visual task state. Similar to the resting state, the PVNH-cortical PLVs were dominant in the α-β band in both MEG (Figure 4F) and sEEG (Figure 4H) visual experiments. Together, the visual response properties and cortical coupling characteristics suggested an essential functional role of PVNH.

## Discussion

In the current study, we applied a state-of-art quantitative analysis approach to characterize the brain-wide inter-areal synchronization in patients with PVNH. Results showed that the synchronization between PVNH and the neocortex and within the neocortex identically predominated in the α-β band during resting and task states. Besides, local neural activities in sensory areas were associated with their coupling strength with the PVNH tissue. Furthermore, in a rare opportunity, we revealed that the PVNH tissue was coactivated with the ipsilateral visual cortex during visual tasks. These results strongly indicate that the PVNH tissue is functionally integrated into neocortical circuits.

Inter-areal coupling at different frequency bands is a general characteristic of the cortical network and is suggested to be an underlying mechanism of neuronal communications in both healthy^18,23,24^ and epileptic^25^ brains. Although previous fMRI studies have revealed the functional connectivity between the PVNH tissue and surrounding^7,9,13–15,26,27^ or distant^9,13^ cortical areas, the oscillatory features of this connectivity are largely unknown. Using whole-brain MEG with high resolution in spectral, temporal, and spatial domains, the current study has found two prominent characteristics of the PVNH-cortical coupling in the resting state, which indicate that the PVNH tissue is functionally integrated into neocortical circuits. One is that PVNH-cortical and cortico-cortical PLVs share a similar spectral feature, a peak in the α-β band, which is similar to the spectral property of cortico-cortical PLVs in healthy participants.^18,23,28^ The other feature is the associated changes between the PVNH-cortical coupling and local neural activities in sensory areas, which has been found in normal cortico-cortical interactions.^29,30^ In addition, the PVNH-cortical coupling strength is not linearly correlated to distance. Therefore, our results reveal that PVNH tissue may be coupled with both anatomically near and distant areas of the neocortex via a shared frequency band.

PVNH has long been considered to be related to the pathological brain network which leads to seizures. Previous studies verified the involvement of the PVNH tissue in epileptic networks with the coactivation of the PVNH tissue and seizure onset zones.^11,31^ However, a recent perspective suggested that there may exist two PVNH-cortical networks, where epileptic tissues in PVNH are connected to epileptic cortical areas,^7,32^ while non-epileptic tissues in PVNH are connected to cognitive cortical networks.^17^ Using sEEG recordings, it was shown that only a portion of recording sites within PVNH tissues generated normal physiological responses and connected to the neocortex (i.e. non-epileptic tissues) during a cognitive task.^17^ In the current study, we further confirm their findings by revealing that the PVNH tissue shows visual evoked responses as well as inter-areal coupling with neocortex during the visual task state.

It is not surprising that the PVNH tissue exhibits visual evoked response which mimics the response in visual cortex ipsilateral to PVNH, in both MEG and sEEG visual experiments. A previous case study found that direct stimulation of the PVNH tissue led to auditory or visual hallucinations,^16^ suggesting a cognitive role of PVNH neurons. This assumption is supported by subsequent fMRI and sEEG studies which reveal that the PVNH tissue is activated during multiple cognitive tasks.^12,17^ In the current study, we additionally probed the relationship between the activities in the PVNH tissue and visual cortex by measuring the peak latencies. Interestingly, no significant difference between the two response latencies was found, in neither the MEG nor the sEEG visual experiment. One possible explanation is that both the PVNH tissue and visual cortex have matured visual neuron assembly to receive parallel synaptic projections from subcortical structures. That is, although neurons in the PVNH tissue are failed to migrate properly, they still have opportunities to develop, mature, and eventually get involved in functional cortical circuits.

Our study has several limitations. First, due to the limited spatial resolution of MEG source localization, we are not able to dissociate MEG signals of epileptic PVNH tissues from those of non-epileptic PVNH tissues. Second, we did not observe any significant phase coupling at high frequencies, which may be restricted by our MEG sampling rate (1000 Hz).^18^ Third, as the opportunity to collect these data is extremely rare, we only inspected the visual function in the PVNH tissue, while other cognitive functions may also relate to the PVNH tissue.^17^ These issues should be investigated with advanced approaches in future studies.

In summary, combining MEG and sEEG recordings, we extend our knowledge that ectopic neurons in the PVNH tissue may retain cognitive functions and be functionally integrated into neocortical circuits, suggesting a potential role for the abnormally migrating neurons in co-development with normally migrating neurons during brain development.

## Acknowledgement

We thank Dr. Yifei Zhang for his constructive comments on the manuscript.

